# Varicose veins of lower extremities: insights from the first large-scale genetic study

**DOI:** 10.1101/368365

**Authors:** Alexandra S. Shadrina, Sodbo Z. Sharapov, Tatiana I. Shashkova, Yakov A. Tsepilov

## Abstract

Varicose veins of lower extremities (VVs) are a common multifactorial vascular disease. Genetic factors underlying VVs development remain largely unknown. Here we report the first large-scale study of VVs performed on a freely available genetic data of 408,455 European-ancestry individuals. We identified 7 reliably associated loci that explain 10% of the SNP-based heritability, and prioritized the most likely causal genes *CASZ1, PPP3R1, EBF1, STIM2*, and *HFE*. Genetic correlation analysis confirmed known epidemiological associations and found genetic overlap with various traits including fluid intelligence score, educational attainment, smoking, and pain. Finally, we observed causal effects of height, weight, both fat and fat-free mass, and plasma levels of MICB and CD209 proteins.

---

Varicose veins (VVs) are one of the clinical manifestations of chronic venous disease posing both a cosmetic and medical problem. VVs can be found in different parts of the body, but most commonly occur in the lower extremities. Prevalence estimates of this condition vary across ethnic groups ranging from 2–4% in the Northern group of the Cook Islands to 50–60% in some countries of the Western world^1^. Increased age, female sex, number of pregnancies, obesity, history of deep venous thrombosis, and standing occupation are among other risk factors^2^. VVs not related to the post-thrombotic syndrome or venous malformations are defined as primary VVs.

Pathogenesis of VVs is still not fully clarified. According to current understanding, key factors implicated in VVs development include changes in hemodynamic forces (decrease in laminar shear stress and increase in venous filling pressure), endothelial activation, inflammation, hypoxia, and dysregulation of matrix metalloproteinases and their tissue inhibitors^3^-^5^. These alterations underlie pathological remodeling of the vascular wall and loss of its tone. Questions remain about the order of events and the primary stimulus triggering the set of disease-related changes.

The cumulative evidence from epidemiological, family, and genetic association studies strongly indicates that there is a hereditary component in VVs etiology^6^-^8^. However, despite progress in this field^9-12^, current knowledge of the genetic basis of this pathology is far from being complete. Elucidating genes involved in susceptibility to VVs would help to identify key molecular players in the disease initiation, provide deeper insights into its pathogenesis, and eventually contribute to development of improved targeted therapy aimed at VVs treating and preventing.

Large-scale biobanks linked to electronic health records open up unparalleled opportunities to investigate the genetics of complex traits. Today, UK Biobank is the largest repository that contains information on genotypes and phenotypes for half a million participating individuals^13^.

This resource is open to all bona fide researchers, and access to data is provided upon approval of their application and payment of necessary costs. However, the need to incur high costs related to data access and computation can be an insurmountable obstacle for those who cannot afford these expenses. The Neale Lab (http://www.nealelab.is/) and the Gene ATLAS database^14^ (http://geneatlas.roslin.ed.ac.uk/) are two independent projects, which intend to remove this burden by generating genetic association data and sharing them with broader scientific community. These resources provide free open access to “quick-and-dirty” genome-wide association study (GWAS) summary statistics for a wide range of phenotypes measured in the UK Biobank. In our study, we aimed to employ state-of-the-art bioinformatics approaches to extract maximum possible information from these open resources with regard to the genetics of VVs of lower extremities. Our objectives were to (1) identify genetic loci reliably associated with VVs risk and prioritize the genes that account for the revealed associations, (2) elucidate pleiotropic effects of identified loci, (3) investigate genetic overlap between VVs and other complex traits, (4) gain etiological insights and explore cause-and-effect relationships by means of Mendelian randomization analysis.

## Results

### Study design, assumptions, and limitations

Our study was designed as a classical GWAS involving the initial discovery screening followed by the replication stage. All calculations were entirely based on the UK Biobank data for white British individuals available in open access databases. The overall workflow of the study is depicted in Fig. 1. In the discovery stage, we used GWAS summary statistics provided by the Neale Lab. To be able to perform a replication, we extracted the UK Biobank data not covered by this project by means of reverse meta-analysis of two overlapping datasets: genetic association data provided by the Gene ATLAS database and the GWAS data supplied by the Neale Lab. The projects used different software and methods of analysis. The Gene ATLAS project^14^ applied less stringent filtering criteria and therefore had larger sample size. This difference in quality control approaches enabled us to generate a subset of “subtracted” individuals and use them as a replication cohort. Data obtained in both GWAS stages were used for further downstream analyses.

**Figure 1.**
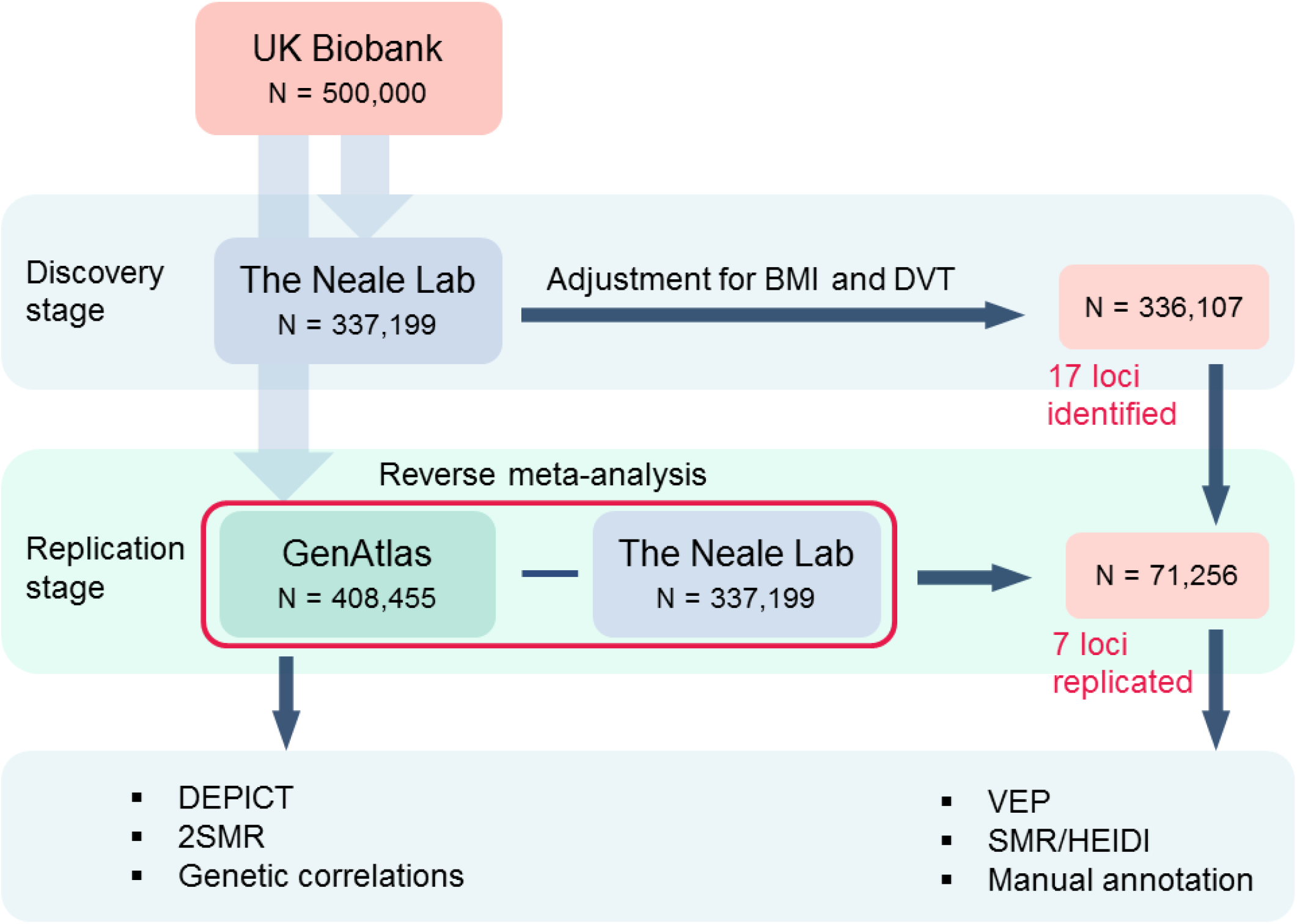
Scheme depicting the overall workflow of our study. In the discovery stage, we used GWAS summary statistics for VVs provided by the Neale Lab and adjusted it for two potential confounders – body mass index (BMI) and deep venous thrombosis (DVT). GWAS data for these traits was also obtained from the Neale Lab database. In order to validate the revealed association signals, we generated a replication cohort by means of reverse meta-analysis using data from by the Gene ATLAS (larger sample size) and the Neale Lab (smaller sample size) databases. VVs traits in all datasets including our simulated replication sample were highly genetically correlated with each other (Supplementary table 1). For 7 replicated loci, we performed a functional annotation analysis. Other analyses were conducted using the Gene ATLAS data since it enables to achieve the highest statistical power and is genetically equivalent to our discovery dataset (rg = 1.00).

In carrying out this study, we had to face a number of challenges and limitations that must be acknowledged. The first limitation was intrinsic to the general approach to phenotype definition based on the electronic medical records system, which was employed in the UK Biobank study. Phenotype “VVs of lower extremities” was defined based on International Classification of Disease (ICD-10) billing code “I83” present in the electronic patient record. The Neale Lab reported the phenotype prevalence of 2.1%. It is much lower than VVs prevalence estimated by European epidemiological studies. Despite the evidence of a “healthy volunteer” selection bias in the UK Biobank study^15^, such a low rate indicates that a proportion of individuals remained undiagnosed. This is in line with a recent primary healthcare register-based study reporting VVs prevalence rate of 3% in German general practice^16^. This phenomenon could be explained by a non-life-threatening nature of varicose veins, which might discourage patients from communication to the doctor. Given that individuals not diagnosed with I83 served as controls in our study, we could therefore expect an overall decrease in the statistical power to detect gene-disease associations. Another potential source of missing associations was immensely strict criteria used by the Neale Lab for single nucleotide polymorphisms (SNPs) quality control removing around 75% of SNPs initially provided by UK Biobank.

The next important limitation arose from the lack of access to individual-level data resulting in the inability to control a possible selection or sampling bias. We suggested that traits related to VVs risk factors could potentially cause unequal representation of patients with different characteristics in the case and the control groups, and thereby induce spurious associations or effect modification. The Neale Lab analyses were adjusted for sex, but other factors were beyond our control. In order to address this challenge, we performed an adjustment for two potential confounders – body mass index (BMI) and deep venous thrombosis (DVT) – by implementation of the method based on GWAS summary statistics^17,18^ (Supplementary Methods).

Last but not least, a noteworthy debatable point in our study is the approach to generating a replication cohort. Due to our intention to use only free open resources, we had to bypass the need to refer to individual patient data. Our replication study sample comprised individuals who have passed the Gene ATLAS quality control criteria, but have been filtered out by the Neale Lab. It was almost 5 times smaller than our discovery study sample. However, although this “dirty” design limited our power to replicate associations, it still could not produce any false positive findings. Moreover, we calculated genetic correlations between VVs in the replication cohort and VVs in both public databases to ensure that these traits are genetically similar (Supplementary table 1). VVs in our replication study was highly correlated with the same trait in other datasets (rg ∼ 0.9). Furthermore, it is noteworthy that VVs in the Gene ATLAS, the Neale Lab as well as in BMI-and DVT-adjusted Neale Lab GWAS were almost genetically equivalent (rg = 0.99 or rg = 1.00).

Summarizing the above, we can state that the limitations which we could not circumvent were mainly related to the loss of statistical power. Nevertheless, given a large sample size and a large number of associations tested, we assume that this obstacle could be at least partially compensated by a huge scale of the UK Biobank study itself.

### Genome-wide association study for VVs of lower extremities

We defined associated loci as regions within 250 kb from the lead SNP and reported only the most significant SNP hits per locus.

In the discovery stage, we identified 18 loci that met a genome-wide level of statistical significance of *P* < 5.0e-08 as provided by the Neale Lab (Table 1, Supplementary table 2). Adjustment for DVT and BMI only slightly changed the observed effects of SNPs, but yielded two additional genome-wide significant loci tagged by rs11693897 and rs247749. However, both these SNPs as well as polymorphism rs1061539 turned insignificant after the correction for the genomic inflation factor (LD Score regression intercept) λ = 1.0403. In the replication stage, 7 loci were replicated at *P* < 0.0029 (0.05/17). Manhattan plot of −log_10_(*P*) is presented in Fig. 2. A quantile-quantile plot for observed vs. expected distribution of *P*-values is shown in Supplementary fig. 2.

**Table 1.** Top SNPs associated with varicose veins of lower extremities. Replicated associations are shown in bold.

| SNP | Chr: position* | EffA/RefA<sup>†</sup> | EAF | Discovery stage |  |  |  |  |  |  | Replication stage,<br>N = 71,256 |  | h<sup>2</sup> (%)<sup>§</sup> |
| --- | --- | --- | --- | --- | --- | --- | --- | --- | --- | --- | --- | --- | --- |
|  |  |  |  | Without adjustment for BMI and DVT,<br>N = 337,199 |  |  | Adjusted for BMI and DVT,<br>N = 336,107 |  |  |  |  |  |  |
| | | | | $\beta$ | SE | $P$ | $\beta$ | SE | $P$ | $P^*_{(GC)}$ | $Z$ | $P$ | |
| **rs11121615** | 1: 10825577 | T/C | 0.69 | -0.0056 | 0.0004 | 1.1e-49 | -0.0056 | 0.0004 | 7.6e-50 | 6.0e-48 | -8.55 | **1.2e-17** | 3.7 |
| **rs2911463** | 16: 88835545 | A/G | 0.69 | -0.0041 | 0.0004 | 1.0e-27 | -0.0041 | 0.0004 | 9.1e-28 | 9.7e-27 | -4.05 | **5.1e-05** | 1.6 |
| **rs2861819** | 2: 68489221 | C/G | 0.66 | 0.0030 | 0.0004 | 8.5e-17 | 0.0030 | 0.0004 | 7.4e-17 | 3.0e-16 | 5.34 | **9.5e-08** | 1.2 |
| **rs3101725** | 5: 127524018 | C/T | 0.76 | -0.0030 | 0.0004 | 1.5e-13 | -0.0030 | 0.0004 | 2.9e-13 | 8.4e-13 | -4.02 | **5.7e-05** | 0.9 |
| **rs11135046** | 5: 158230013 | T/G | 0.54 | -0.0025 | 0.0003 | 5.9e-13 | -0.0025 | 0.0003 | 3.7e-13 | 1.1e-12 | -4.13 | **3.6e-05** | 0.9 |
| **rs28558138** | 4: 26818080 | C/G | 0.42 | -0.0024 | 0.0004 | 1.8e-11 | -0.0024 | 0.0004 | 1.3e-11 | 3.2e-11 | -6.39 | **1.7e-10** | 1.1 |
| **rs7773004** | 6: 26267755 | G/A | 0.49 | -0.0023 | 0.0003 | 5.2e-11 | -0.0023 | 0.0003 | 6.4e-11 | 1.5e-10 | -3.05 | **0.002** | 0.6 |
| rs7856039 | 9: 118385117 | C/T | 0.67 | 0.0024 | 0.0004 | 5.7e-11 | 0.0024 | 0.0004 | 8.1e-11 | 1.9e-10 | 0.10 | 0.92 | – |
| rs9880192 | 3: 128297569 | C/G | 0.41 | -0.0023 | 0.0004 | 8.0e-11 | -0.0023 | 0.0004 | 8.7e-11 | 2.0e-10 | -2.03 | 0.04 | – |
| rs192647746 | 4: 182760366 | G/A | 0.001 | 0.0336 | 0.0054 | 4.5e-10 | 0.0333 | 0.0052 | 1.2e-10 | 2.9e-10 | -1.01 | 0.31 | – |
| rs12625547 | 20: 50154647 | G/T | 0.17 | -0.0029 | 0.0005 | 1.2e-10 | -0.0029 | 0.0005 | 1.3e-10 | 2.9e-10 | -2.18 | 0.03 | – |
| rs236530 | 17: 68217471 | T/C | 0.75 | -0.0026 | 0.0004 | 1.8e-10 | -0.0026 | 0.0004 | 1.4e-10 | 3.2e-10 | -2.84 | 0.005 | – |
| rs2241173 | 17: 70028445 | G/A | 0.58 | -0.0022 | 0.0004 | 4.5e-10 | -0.0022 | 0.0004 | 4.4e-10 | 9.7e-10 | -1.84 | 0.07 | – |
| rs73107980 | 12: 48187601 | T/C | 0.24 | 0.0025 | 0.0004 | 1.7e-09 | 0.0025 | 0.0004 | 1.4e-09 | 2.8e-09 | 1.65 | 0.10 | – |
| rs10084824 | 4: 89515141 | A/C | 0.01 | 0.0100 | 0.0018 | 4.8e-08 | 0.0101 | 0.0018 | 1.2e-08 | 2.3e-08 | -1.75 | 0.08 | – |
| rs6062619 | 20: 62683002 | G/A | 0.27 | -0.0022 | 0.0004 | 2.0e-08 | -0.0022 | 0.0004 | 1.4e-08 | 2.6e-08 | -2.80 | 0.005 | – |
| rs1936800 | 6: 127436064 | T/C | 0.53 | -0.0019 | 0.0003 | 2.0e-08 | -0.0020 | 0.0003 | 1.9e-08 | 3.6e-08 | -0.31 | 0.76 | – |
| rs11693897 | 2: 45933993 | T/C | 0.004 | 0.0148 | 0.0029 | 3.3e-07 | 0.0149 | 0.0027 | 3.3e-08 | 6.2e-08 | -0.94 | 0.34 | – |
| rs247749 | 5: 76296735 | C/T | 0.31 | 0.0021 | 0.0004 | 5.5e-07 | 0.0021 | 0.0004 | 3.9e-08 | 7.3e-08 | 1.78 | 0.07 | – |
| rs1061539 | 6: 29937555 | C/T | 0.83 | -0.0025 | 0.0005 | 4.0e-08 | -0.0025 | 0.0005 | 4.8e-08 | 8.9e-08 | -2.69 | 0.007 | – |

BMI, body mass index; DVT, deep venous thrombosis; EAF, effect allele frequency; SE, standard error; SNP, single nucleotide polymorphism
\*Chromosome: position on chromosome according to GRCh37.p13 assembly
<sup>†</sup>Effect allele/reference allele
<sup>‡</sup>P-value corrected for the genomic inflation factor
<sup>§</sup>Proportion of SNP-based heritability explained by the variant

**Figure 2.**
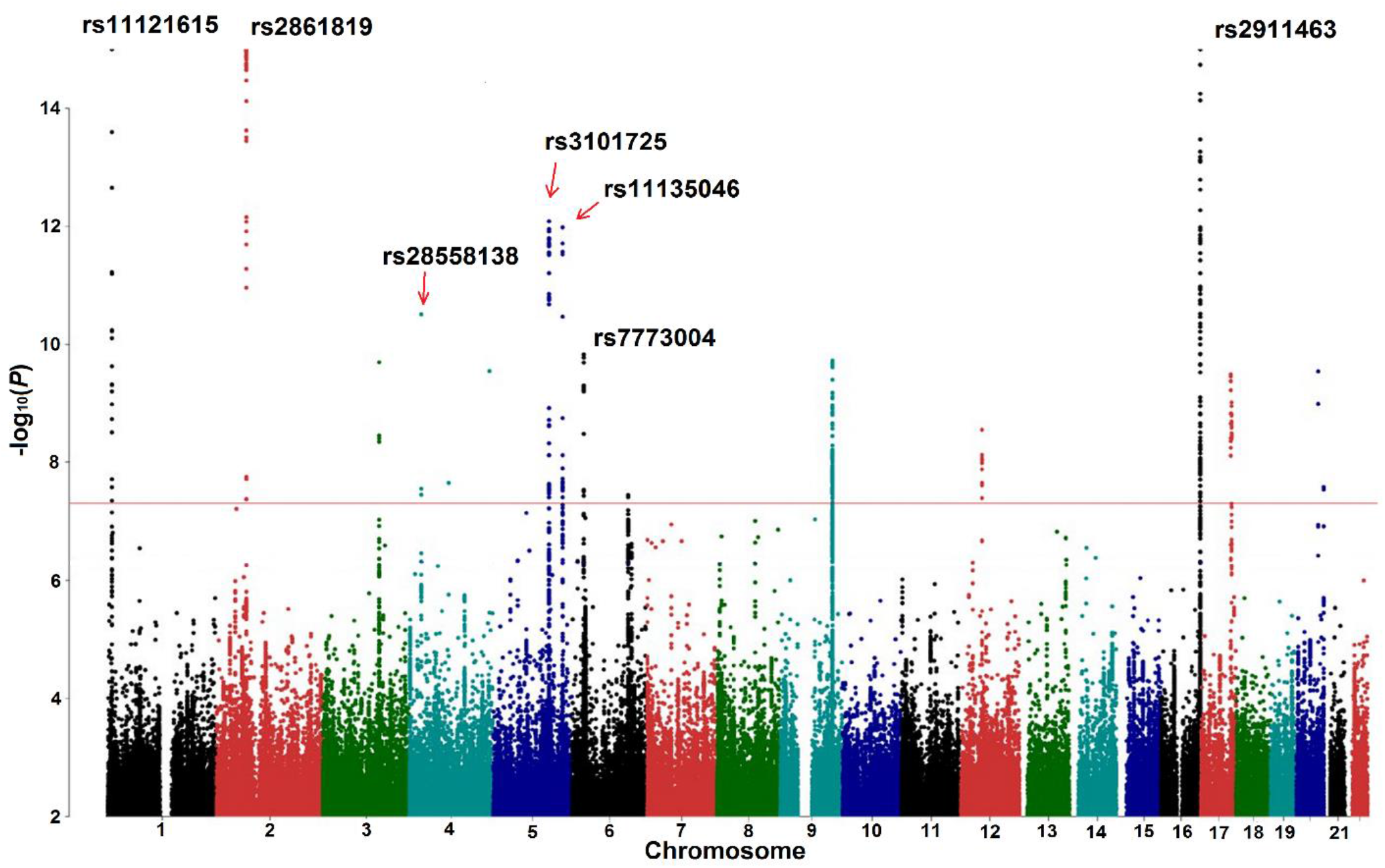
Graphical summary of the discovery GWAS stage after the adjustment for deep venous thrombosis and body mass index and the correction for genomic control. Red line corresponds to the genome-wide significance threshold of *P* = 5.0e-08. Only associations with *P* < 1.0e-02 are presented. Points with −log_10_(*P*) >15 are depicted as points with −log_10_(*P*) =15.

Regional association plots for replicated loci are given in Supplementary fig. 1. Genes located in these regions are listed in Supplementary table 2. We performed conditional and joint (COJO) analysis and detected two independent signals in the locus on chromosome 16 at a distance of 39 kb from each other (Supplementary table 3). Lead SNPs rs9972645 and rs2911463 were in low linkage disequilibrium (r^2^ = 0.15).

### Functional annotation of the revealed signals

#### Literature-based annotation

We explored whether genes in close proximity to replicated hits could have biologically plausible roles in VVs based on their functions or previously revealed associations with other complex traits. The results of literature-based candidate gene prioritization are provided in Table 2. Furthermore, we compiled a list of traits associated at a genome-wide significance level with either lead SNPs, or variants in linkage disequilibrium (LD) with these polymorphisms using both the Pubmed and the PhenoScanner databases (Table 2, Supplementary table 4). Our analysis showed that among the genes nearest to the top GWAS loci (±250 kb), 2 genes were related to vascular development and remodeling (*CASZ1* and *PIEZO1*) with one gene having putative role in vascular remodeling (*STIM2*), 4 genes were implicated in blood pressure and hypertension (*CASZ1, PIEZO1, SLC12A2*, and *EBF1*), 2 genes were linked to inflammation (*PPP3R1* and *EBF1*), and 2 genes were ion channels involved in regulation of both cell volume and vascular tone (*PIEZO1* and *SLC12A2*). Near the *SLC12A2*, we identified the *FBN2* gene which encodes a protein regulating elastogenesis, so we prioritized both genes.

**Table 2.**
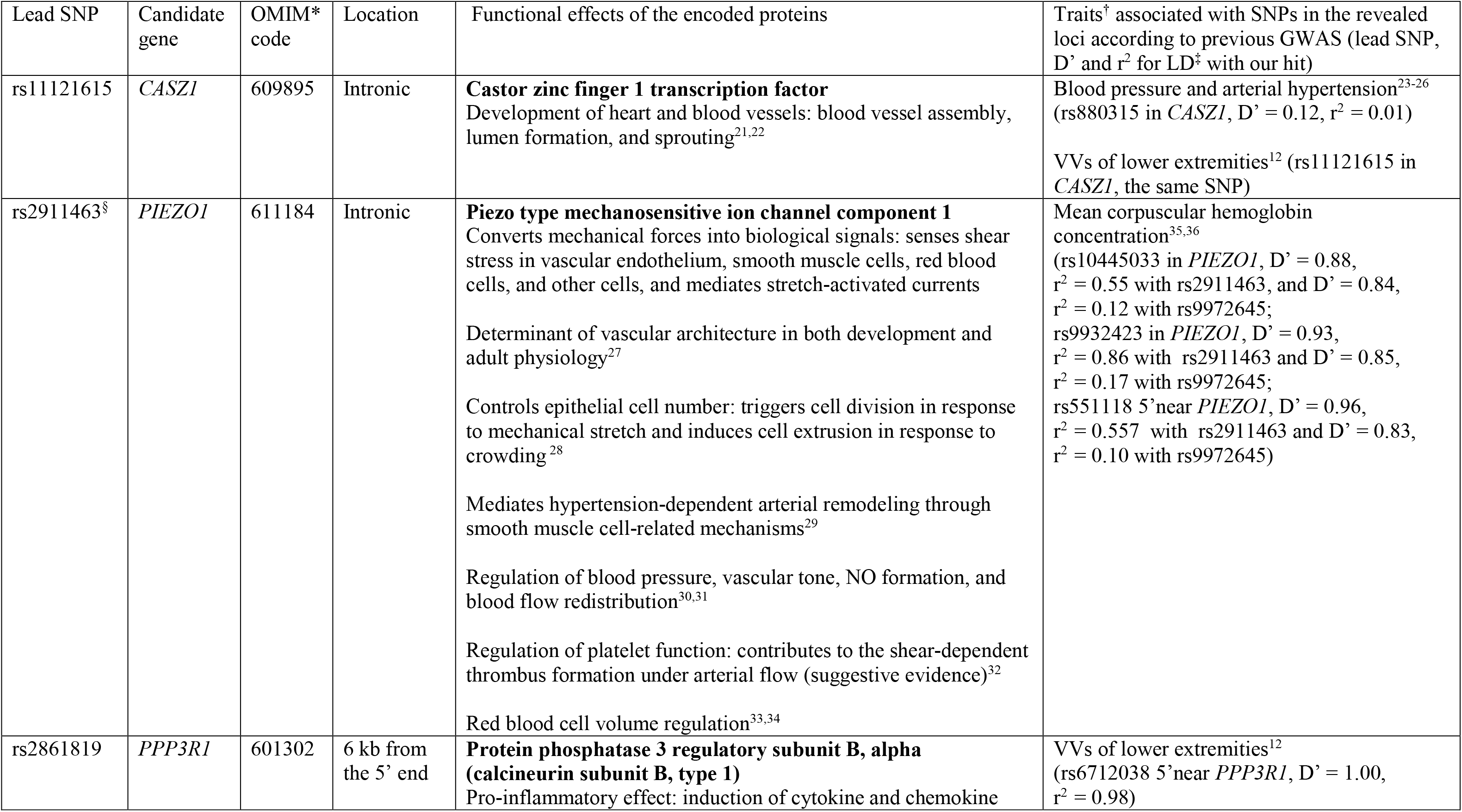
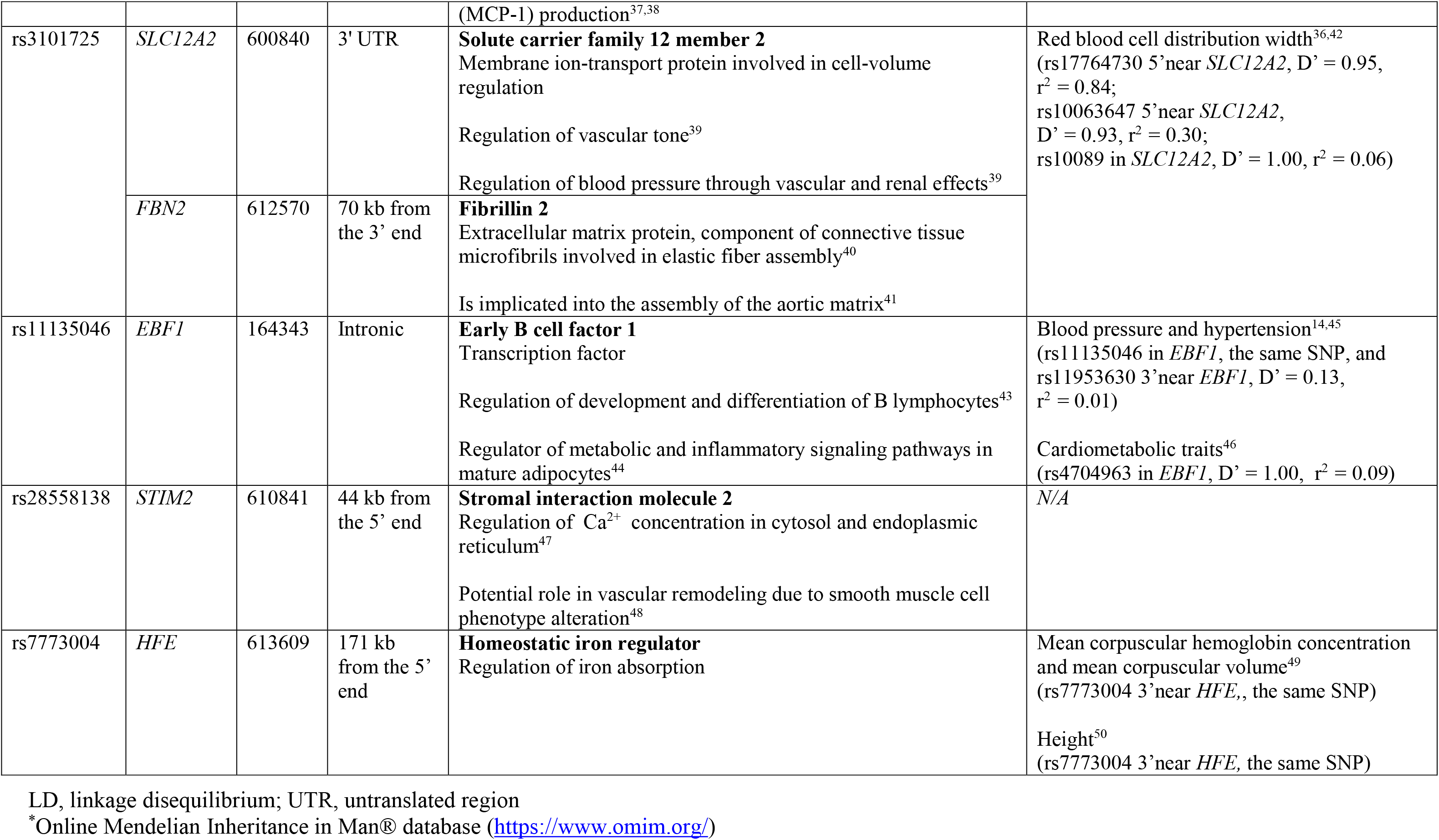
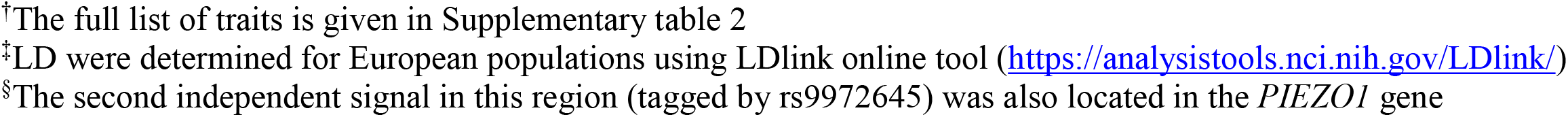
Gene prioritization based on a literature review.

| Lead SNP | Candidate gene | OMIM* code | Location | Functional effects of the encoded proteins | Traits<sup>†</sup> associated with SNPs in the revealed loci according to previous GWAS (lead SNP, D' and r<sup>2</sup> for LD<sup>‡</sup> with our hit) |
| --- | --- | --- | --- | --- | --- |
| rs11121615 | *CASZ1* | 609895 | Intronic | **Castor zinc finger 1 transcription factor**<br>Development of heart and blood vessels: blood vessel assembly, lumen formation, and sprouting<sup>21,22</sup> | Blood pressure and arterial hypertension<sup>23-26</sup><br>(rs880315 in *CASZ1*, D' = 0.12, r<sup>2</sup> = 0.01)<br>VVs of lower extremities<sup>12</sup> (rs11121615 in *CASZ1*, the same SNP) |
| rs2911463<sup>§</sup> | *PIEZO1* | 611184 | Intronic | **Piezo type mechanosensitive ion channel component 1**<br>Converts mechanical forces into biological signals: senses shear stress in vascular endothelium, smooth muscle cells, red blood cells, and other cells, and mediates stretch-activated currents<br>Determinant of vascular architecture in both development and adult physiology<sup>27</sup><br>Controls epithelial cell number: triggers cell division in response to mechanical stretch and induces cell extrusion in response to crowding<sup>28</sup><br>Mediates hypertension-dependent arterial remodeling through smooth muscle cell-related mechanisms<sup>29</sup><br>Regulation of blood pressure, vascular tone, NO formation, and blood flow redistribution<sup>30,31</sup><br>Regulation of platelet function: contributes to the shear-dependent thrombus formation under arterial flow (suggestive evidence)<sup>32</sup><br>Red blood cell volume regulation<sup>33,34</sup> | Mean corpuscular hemoglobin concentration<sup>35,36</sup><br>(rs10445033 in *PIEZO1*, D' = 0.88, r<sup>2</sup> = 0.55 with rs2911463, and D' = 0.84, r<sup>2</sup> = 0.12 with rs9972645;<br>rs9932423 in *PIEZO1*, D' = 0.93, r<sup>2</sup> = 0.86 with rs2911463 and D' = 0.85, r<sup>2</sup> = 0.17 with rs9972645;<br>rs551118 5' near *PIEZO1*, D' = 0.96, r<sup>2</sup> = 0.557 with rs2911463 and D' = 0.83, r<sup>2</sup> = 0.10 with rs9972645) |
| rs2861819 | *PPP3R1* | 601302 | 6 kb from the 5' end | **Protein phosphatase 3 regulatory subunit B, alpha (calcineurin subunit B, type 1)**<br>Pro-inflammatory effect: induction of cytokine and chemokine | VVs of lower extremities<sup>12</sup><br>(rs6712038 5' near *PPP3R1*, D' = 1.00, r<sup>2</sup> = 0.98) |
|  |  |  |  | (MCP-1) production<sup>37,38</sup> |  |
| rs3101725 | *SLC12A2* | 600840 | 3' UTR | **Solute carrier family 12 member 2**<br>Membrane ion-transport protein involved in cell-volume regulation<br>Regulation of vascular tone<sup>39</sup><br>Regulation of blood pressure through vascular and renal effects<sup>39</sup> | Red blood cell distribution width<sup>36,42</sup><br>(rs17764730 5' near *SLC12A2*, $D' = 0.95$, $r^2 = 0.84$;<br>rs10063647 5' near *SLC12A2*, $D' = 0.93$, $r^2 = 0.30$;<br>rs10089 in *SLC12A2*, $D' = 1.00$, $r^2 = 0.06$) |
|  | *FBN2* | 612570 | 70 kb from the 3' end | **Fibrillin 2**<br>Extracellular matrix protein, component of connective tissue microfibrils involved in elastic fiber assembly<sup>40</sup><br>Is implicated into the assembly of the aortic matrix<sup>41</sup> |  |
| rs11135046 | *EBF1* | 164343 | Intronic | **Early B cell factor 1**<br>Transcription factor<br>Regulation of development and differentiation of B lymphocytes<sup>43</sup><br>Regulator of metabolic and inflammatory signaling pathways in mature adipocytes<sup>44</sup> | Blood pressure and hypertension<sup>14,45</sup><br>(rs11135046 in *EBF1*, the same SNP, and rs11953630 3' near *EBF1*, $D' = 0.13$, $r^2 = 0.01$)<br>Cardiometabolic traits<sup>46</sup><br>(rs4704963 in *EBF1*, $D' = 1.00$, $r^2 = 0.09$) |
| rs28558138 | *STIM2* | 610841 | 44 kb from the 5' end | **Stromal interaction molecule 2**<br>Regulation of $Ca^{2+}$ concentration in cytosol and endoplasmic reticulum<sup>47</sup><br>Potential role in vascular remodeling due to smooth muscle cell phenotype alteration<sup>48</sup> | N/A |
| rs7773004 | *HFE* | 613609 | 171 kb from the 5' end | **Homeostatic iron regulator**<br>Regulation of iron absorption | Mean corpuscular hemoglobin concentration and mean corpuscular volume<sup>49</sup><br>(rs7773004 3' near *HFE*, the same SNP)<br>Height<sup>50</sup><br>(rs7773004 3' near *HFE*, the same SNP) |

LD, linkage disequilibrium; UTR, untranslated region
\*Online Mendelian Inheritance in Man® database (https://www.omim.org/)
†The full list of traits is given in Supplementary table 2
‡LD were determined for European populations using LDlink online tool (https://analysistools.nci.nih.gov/LDlink/)
§The second independent signal in this region (tagged by rs9972645) was also located in the *PIEZO1* gene

Two signals overlapped with findings from a recent GWAS for VVs conducted by “23andMe” company^12^: rs11121615 in the *CASZ1* gene, which was among the top SNPs identified in that study, and rs2861819 near the *PPP3R1*, which was in complete LD (D’ = 1.00, r^2^ = 0.98) with their another hit rs6712038. For 3 loci (tagged by rs2911463, rs3101725, and rs7773004) a genome-wide significant association with red blood cell traits has previously been demonstrated. One of these signals was near the *HFE* gene implicated in iron absorption. The same locus showed association with height.

Changes in blood pressure, abnormal vascular wall remodeling, altered vascular tone, deterioration of vein wall elastic properties as well as inflammation are known factors related to VVs. Association with red blood cell traits is intriguing. Hypothetically, both erythrocyte homeostasis and VVs could be linked to iron overload^19,20^. It is noteworthy that association between hemochromatosis-related functional variant in the *HFE* gene and primary VVs was revealed in our previous candidate-gene study in ethnic Russian individuals^19^.

#### Searching for functional variants

Ensembl Variant Effect Predictor (VEP)^51^ analysis identified one moderate-impact missense variant rs2044693 in the *PNO1* gene (in the locus tagged by rs2861819). SIFT and PolyPhen tools predicted no damaging effect of this SNP on the protein function. Additionally, we revealed 19 SNPs in regulatory regions (in loci tagged by rs9972656, rs2861819, rs11135046, and rs7773004), and one synonymous variant rs1368298 in the *EBF1* gene (in the locus tagged by rs11135046). The full data is available in Supplementary table 5.

#### Testing for pleiotropy: VVs and gene expression

We used Summary data-based Mendelian Randomization (SMR) analysis followed by the Heterogeneity in Dependent Instruments (HEIDI)^52^ test to identify genes whose expression level is associated with the same causal SNPs that affect the risk of VVs. The results are given in Supplementary table 6. SMR/HEIDI analysis provided evidence for pleiotropic effects of variants within rs3101725 and rs2861819 regions. In the first region, association was revealed with the expression of long intergenic non-protein coding RNA 1184 (*LINC01184*) located in close proximity to the *SLC12A2* on the reverse strand. Changes in the *SLC12A2* expression were not associated with VVs in our tests. Interestingly, the previous study showed that disruption of a bi-directional promoter between these genes altered the expression of both *SLC12A2* and *LINC01184*, while deletion of the third exon of *LINC01184* affected only *LINC01184* expression^36^.

In the locus tagged by rs2861819, our analysis identified four genes – *PPP3R1, PLEK, WDR92*, and *PNO1*. It is likely that this region contains a strong eQTL that affects the expression of nearby genes in different tissues and influences the risk of VVs.

It is important to note that SMR/HEIDI analysis does not distinguish pleiotropy from causality. Therefore, we can hypothesize causal relationships between the expression level of identified genes and the risk of VVs development.

Besides this, we found associations with expression levels of four genes within rs2911463 region (*APRT, ZFPM1, RNF166*, and *PIEZO1*) as well as with expression level of the *TRIM38* gene located ∼280 kb from rs7773004. However, statistically significant heterogeneity revealed in the HEIDI test indicated that these associations were mediated by polymorphisms different from those that alter VVs risk.

#### DEPICT analysis

Data-driven Expression Prioritized Integration for Complex Traits (DEPICT) framework^53^ was used to conduct a gene set and tissue enrichment analysis as well as to provide additional evidence for gene prioritization. Results of DEPICT analysis for SNP sets associated with VVs at *P* < 5.0e-08 and *P* < 1.0e-05 are given in Supplementary tables 7 and 8, respectively. For the first set, we identified a significant enrichment of the Mammalian Phenotype (MP) Ontology terms “abnormal vasculogenesis” and “abnormal vascular development”. When we relaxed significance threshold of input SNPs to *P* < 1.0e-05, 22 MP terms were shown to be enriched with additional 9 categories related to abnormal morphology of heart and vessels (“abnormal blood vessel morphology”, “abnormal vascular smooth muscle morphology”, “failure of heart looping”, etc.). The remaining MP terms were associated with aberrant morphology of other tissues and organs and disruption of embryonic development. Besides this, we observed enrichment for several subnetworks of protein-protein interactions, including that for ENG (major glycoprotein of the vascular endothelium, mutations in its gene cause multisystemic vascular dysplasia), SMAD 2, 4, 6, 7 (transduce signals from TGF-beta family members), and Notch4 (regulates arterial specification^54^).

Tissue enrichment analysis revealed no statistically significant categories.

DEPICT gene prioritization tool provided 2 and 47 prioritized genes depending on the level of statistical significance of the input SNPs (*P* < 5.0e-08 and *P* < 1.0e-05, respectively; Supplementary tables 7 and 8).

#### Summary of gene prioritization

Basing on cumulative evidence from different analyses, we drew up a list of genes which most likely account for the revealed associations and seem to be functionally relevant to VVs development (Table 3). For two loci on chromosomes 2 and 5, we were unable to prioritize any candidate gene. However, for the locus on chromosome 2, we advocate for the *PPP3R1* gene since it displayed the lowest *P* and the greatest effect size in the SMR test (*P =*1.3e-22 and *β* = 120.0, Supplementary table 6). Moreover, biological functions of PPP3R1 suggest its involvement in VVs etiology (Table 2). In particular, it induces MCP-1 production, which was shown to be enhanced in VVs^55^,^56^. Finally, a recent study demonstrated that expression of *PPP3R1* is increased in the venous tissue of VVs patients compared with veins of healthy individuals (fold change = 1.5)^57^.

**Table 3.** Summary of gene prioritization.

| Lead SNP | Locus* | Number of genes per locus<sup>†</sup> | Prioritized gene | Nearest gene, *yes/no* | Evidence for prioritization |
| --- | --- | --- | --- | --- | --- |
| rs11121615 | 1: 11075577–10575577 | 4 | *CASZ1* | *yes* | L, D |
| rs2911463 | 16: 89085545–88585545<sup>‡</sup> | 20 | *PIEZO1* | *yes* | L, D |
| rs2861819 | 2: 68739221–68239221 | 8 | *PPP3R1* | *yes* | L, E, S |
|  |  |  | *PLEK* | *no* | S |
|  |  |  | *WDR92* | *no* | S |
|  |  |  | *PNO1* | *no* | V, S |
| rs3101725 | 5: 127774018–127274018 | 3 | *SLC12A2* | *yes* | L |
|  |  |  | *FBN2* | *no* | L |
|  |  |  | *LINC01184* | *no* | S |
| rs11135046 | 5: 158480013–157980013 | 1 | *EBF1* | *yes* | L |
| rs28558138 | 4: 27068080–26568080 | 2 | *STIM2* | *no* | L, D |
| rs7773004 | 6: 26517755–26017755 | 41 | *HFE* | *no* | L |

D, DEPICT analysis; L, literature-based prioritization; E, expression change in VVs; S, SMR/HEIDI analysis; V, Variant Effect Predictor analysis
\*Chromosome: position on chromosome according to GRCh37.p13 assembly
<sup>†</sup>Calculated based on regional association plots generated with LocusZoom tool (http://locuszoom.org/, Supplementary fig.1)
<sup>‡</sup>This locus contains two independent association signals. SNP rs9972645 that tags the second signal is also located in the *PIEZO1* gene

### Testing for pleiotropy: VVs and other complex traits

SMR/HEIDI analysis revealed 31 traits that shared the same casual variants with VVs (traits for this and further analyses were obtained from the GWAS-MAP database, see Methods section for details). These SNPs were located in 5 out of 7 loci identified in our study (Fig. 3, Supplementary table 10). Variants within loci tagged by rs11121615 and rs3101725 showed positive SMR beta coefficients (the same direction of effect) with predicted and fat-free mass of both legs and a whole body as well as with basal metabolic rate, and negative coefficients (opposed effects) – with leg fat percentage as well as impedance of both legs and a whole body (parameter negatively correlated with fat-free mass). Loci tagged by rs3101725 and rs7773004 comprised SNPs with pleiotropic effects on red blood cell erythrocyte distribution width (negative beta). Locus tagged by rs7773004 was also related to numerous blood traits, such as mean corpuscular haemoglobin concentration, mean platelet thrombocyte volume, and monocyte and reticulocyte count. Three loci (rs11135046, rs28558138, and rs28558138) were linked to blood pressure and hypertension with different SMR beta signs. Another interesting finding was identification of positive SMR beta for associations with vascular/heart problems (rs28558138 locus), cellulitis (rs3101725 locus), forced viral capacity, and height size at age 10 (rs7773004 locus). Overall, our analysis revealed four main groups of traits: one cluster related mainly to mass, one cluster involving mostly fat-and blood pressure-related traits, one cluster of blood-related traits linked only to rs7773004 locus, and one independent trait “red blood cell erythrocyte distribution width” (Fig. 3).

**Figure 3.**
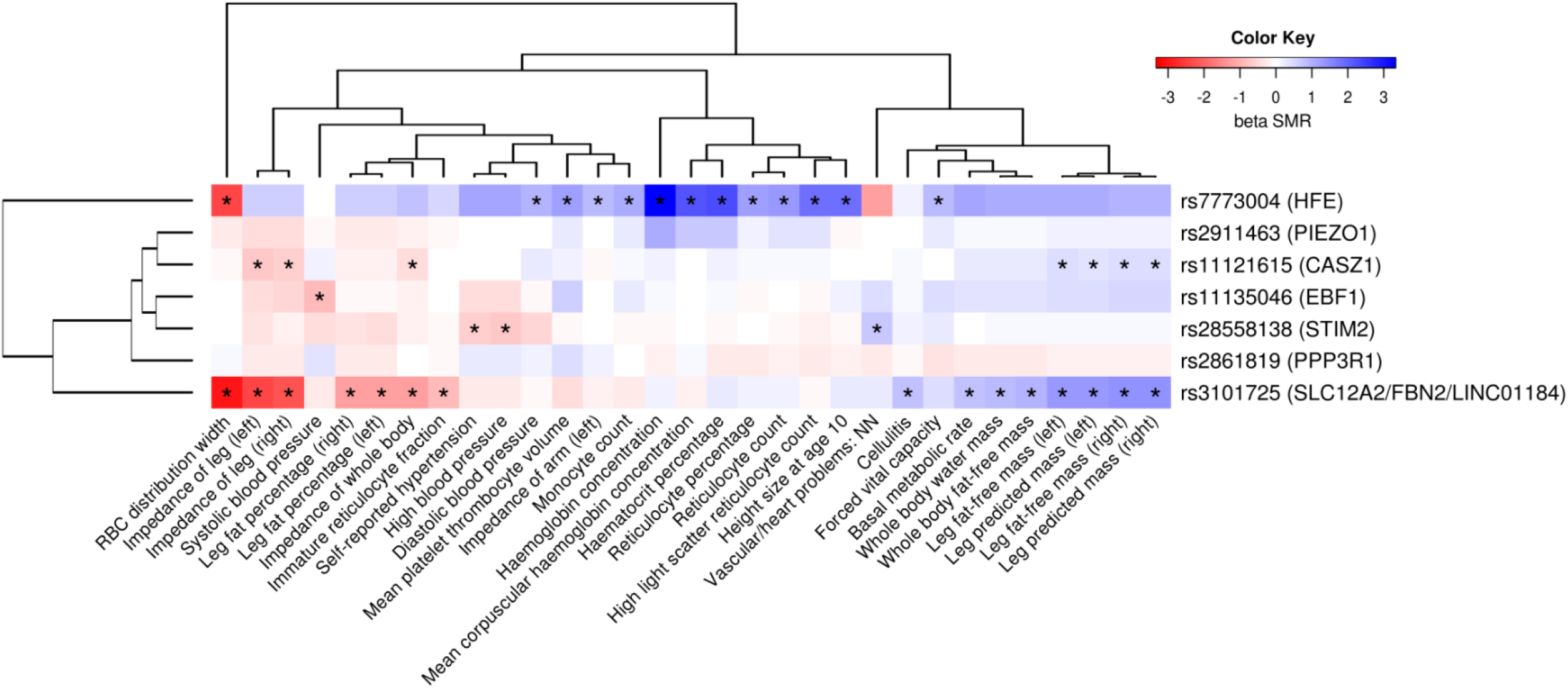
Pleiotropic effects of variants within revealed loci on other complex traits. Color depicts the sign and the magnitude of SMR beta coefficient. Negative sign (red) means opposed effects, and positive sign (blue) means the same direction of effect. Traits that passed both SMR and HEIDI test are marked with an asterisk. Prioritized genes in loci are shown in brackets. Full names of traits as indicated in the Neale lab and the Gene ATLAS databases are given in Supplementary table 14. RBC, red blood cell.

### Genetic correlations

Genetic correlations between VVs in different datasets (discovery and replication cohorts, etc.) are given in Supplementary table 1. A list of genetic correlation estimates (rg) between VVs and 861 complex traits is presented in Supplementary table 11. Twenty five traits showed statistically significant correlation with VVs with absolute values of rg ≥ 0.2. Correlation matrix for this subset is displayed as a heatmap in Fig. 4. We observed 5 main clusters: traits related to the type of job, intelligence, and qualification; traits related to height and mass (including predicted and fat-free mass of both legs); thrombosis-related traits; traits related to operations; and traits related to pain (including leg pain and gonarthrosis) and health satisfaction. VVs trait was closest to the thrombosis-related cluster (positive correlation). It was also positively correlated with mass, operations, and pain-related traits as well as with lower levels of qualification and heavy manual/walking/standing job. For traits with less prominent correlation with VVs, we observed the same trend: pain and anthropometric traits (sitting and standing height, BMI, mass, etc.) showed positive correlations, whilst higher levels of education – negative. Interestingly, negative correlation was observed with usual walking pace, and positive – with current smoking.

**Figure 4.**
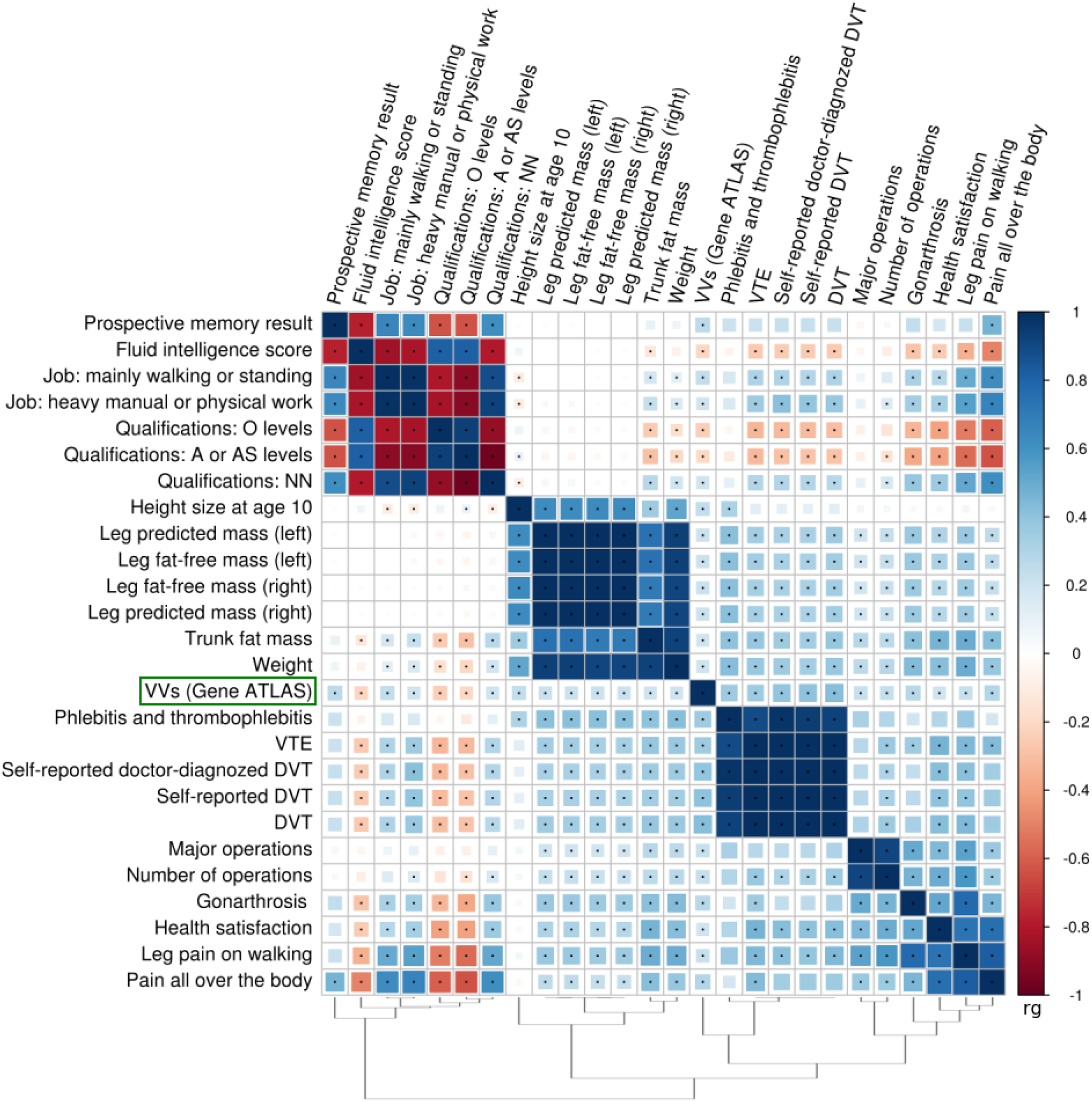
Matrix of genetic correlations between VVs and other complex traits. Color depicts the sign and the absolute value of genetic correlation coefficient (rg). Only the traits having |rg| ≥ 2 with VVs are shown. Combinations with statistically significant rg are marked with points (*P* < 3.1e-05). O levels of qualification correspond to the General Certificate of Education Ordinary Level/the General Certificate of Secondary Education; A or AS level correspond to Advanced or Advanced Subsidiary levels. “Qualifications: NN” means no qualification. Full names of traits as indicated in the Neale lab and the Gene ATLAS databases are given in Supplementary table 15. For “Prospective memory result” and “Health satisfaction”, high scores correspond to poor performance. DVT, deep venous thrombosis; VTE, venous thromboembolism.

We calculated partial genetic correlations for the subset of 7 non-collinear traits with |rg |≥ 0.2. Two traits – DVT and height size at age 10 – were shown to share common genetic background with VVs after the adjustment for the remaining traits in the analyzed subset (Supplementary fig. 3). In other words, their shared genetic components were demonstrated to be at least partially independent of other complex traits. Additionally, we estimated partial genetic correlations between VVs, standing height, and weight. Both standing height and weight had independent genetic components shared with VVs (Supplementary fig. 4).

### Hypothesis-free search for causal relationships

We applied a 2-sample Mendelian randomization (2SMR)^58^ strategy to infer causal relationships between a broad range of “exposure” phenotypes and VVs as an outcome. In total, 39 complex traits were shown to be potential causative factors. Although only genome-wide associated SNPs from exposure GWAS were selected as instrumental variables, we did not require these loci to be replicated. Nevertheless, we checked the stability of our tests with regard to instruments selection by performing the robustness analysis. This test along with the Steiger test^59^ for the correct direction of effect underpinned the exclusion of 2 out of 39 traits (Supplementary table 12). Further, we assessed violations of MR assumption of absence of horizontal pleiotropy (influence of genetic instruments on the outcome only through the exposure, also known as “exclusion-restriction criterion”) by means of sensitivity analyses^58^. We did not observe a statistically significant intercept in MR-Egger regression for any trait. However, only a small proportion of traits showed symmetry in Funnel plots and had no heterogeneity in causal effects amongst instruments. This provides evidence that, for the majority of traits, at least some of the selected instruments exhibit horizontal pleiotropic effects.

Such traits mainly involved several hundred genome-wide SNPs that made leave-one-out analysis also uninformative. In order to correct for horizontal pleiotropy, we applied a straightforward approach having excluded all instrumental variables associated with VVs at the level of statistical significance higher than 0.01. Our correction led to symmetry in 26 out of 33 asymmetrical Funnel plots and eliminated heterogeneity in causal effects for 28 out of 34 traits, although 6 traits lost the statistical significance of 2SMR coefficients (Supplementary fig. 5–42, Supplementary table 12). Removing potential sources of heterogeneity also reduced absolute values of 2SMR beta for all the corrected phenotypes. A graphical representation of our results, including 25 causal inferences that we consider the most reliable, is shown in Fig. 5. Twenty one traits were related to anthropometry and included standing and sitting height, weight, hip and waist circumference, fat-free, predicted, and fat mass of legs and arms, etc. One trait was a spirometry measurement associated with pulmonary function. Nonetheless, since it is positively correlated with height, we suppose that it has no independent effect on VVs development. Similarly, the reverse association between malabsorption/coeliac disease and VVs could actually be induced by a weight loss as one of the complications of these conditions. Moreover, although this trait has passed all the necessary tests, we avoid making strong claims about its causality since it was self-reported and involved only ∼1,500 cases.

**Figure 5.**
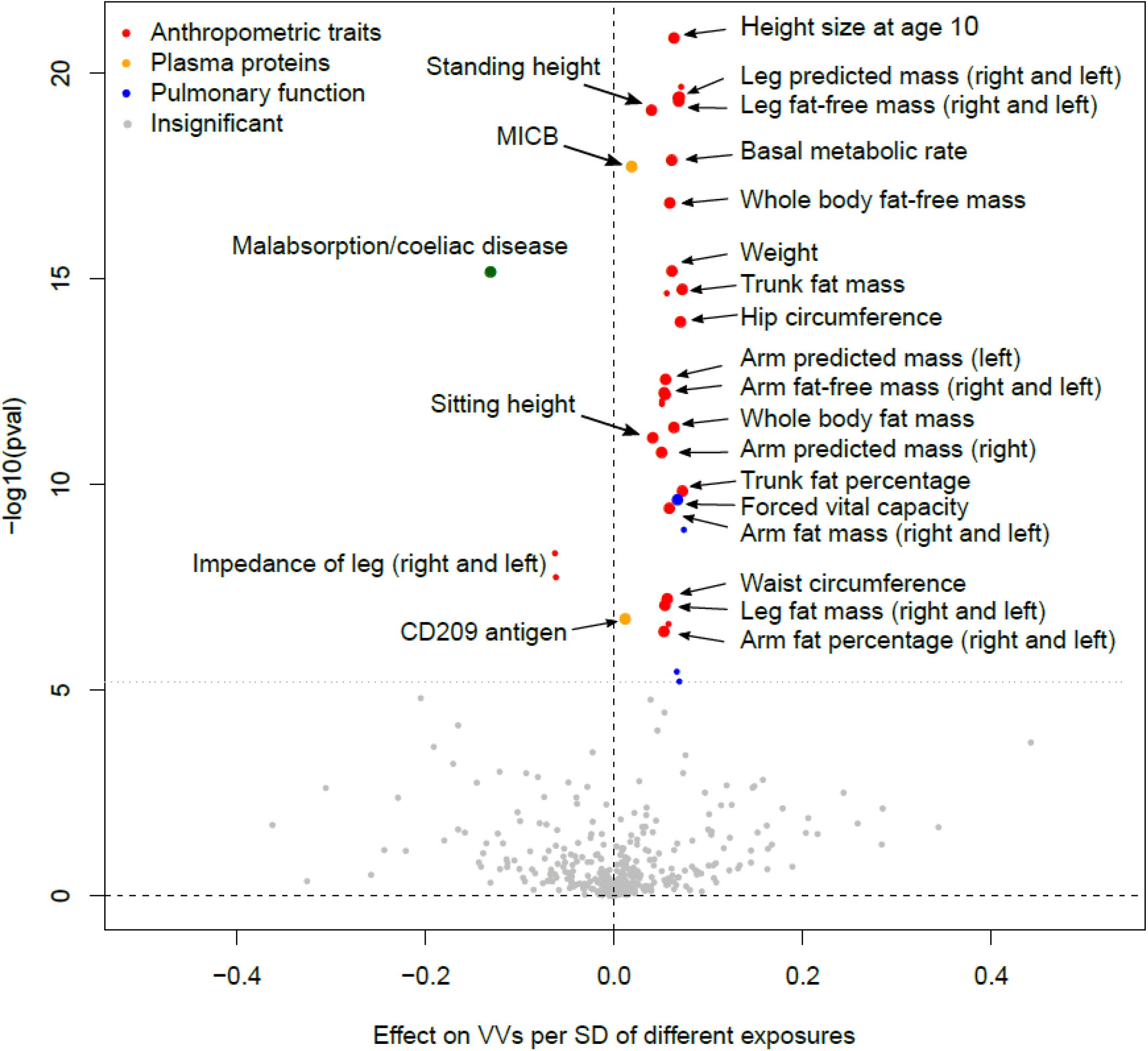
Volcano plot depicting the results of hypothesis-free 2SMR analysis (IVW approach). The x-axis shows 2SMR beta, and the y-axis denotes the level of statistical significance on a logarithmic scale. Grey color indicates the traits that did not pass either 2SMR, or the Steiger test, or additional robustness analysis. Small colored circles depict the traits that passed these test, but either exhibited heterogeneity in causal effects amongst instruments after removing potential outlier SNPs, or did not passed 2SMR after this procedure. Large colored circles represent the traits that both passed 2SMR and showed no heterogeneity after the outliers removal (“the most reliable traits”). 2SMR coefficients are shown as calculated before the correction for pleiotropy. “Pulmonary function” traits include different estimates of forced vital capacity and forced expiratory volume in 1-second. Further details are provided in Supplementary table 12. MICB, MHC class I polypeptide-related sequence B.

All the above mentioned traits were derived from the Neale Lab repository and therefore had about 80% sample overlap with VVs dataset obtained from the Gene ATLAS. Shared participants between the “exposure” and the “outcome” GWAS can cause bias in the Mendelian randomization analysis when weak instruments are selected^60^. In our study, this limitation was partially mitigated by using only strong instruments associated with exposure traits at a high level of statistical significance (*P* < 1.0e-08). In order to fully overcome the problem and confirm our results, we searched for independent GWAS and performed a replication analysis. Data were obtained for height, weight, waist circumference, and body fat percentage. Only height reached Bonferroni-corrected level of statistical significance and remained significant after the correction for pleiotropy (2SMR *P* < 1.0e-07, Supplementary table 13). However, null results for other traits could be explained by a limited power of the analysis. For example, sample size for weight was 4.5 times smaller than in the Neale Lab study.

Finally, the remaining two traits from the “reliable” set were plasma levels of MHC class I polypeptide-related sequence B protein and CD209 antigen. These phenotypes were not UK Biobank traits, therefore our results could not be confounded by a sample overlap.

## Discussion

In the present study, we utilized freely available GWAS summary data to unravel the genetic underpinnings of VVs of lower extremities. Lack of individual-level data made us be as rigorous as possible to avoid false positive discoveries, so we skimmed off the most apparent evidence. Nevertheless, as far as we are aware, our study is the largest and the most comprehensive study of VVs genetics to date. Beyond that, the availability of GWAS results makes our research fully reproducible.

We identified and replicated 7 susceptibility loci that explain 10% of the SNP-based heritability, and prioritized the most likely causal genes (Fig.2, Tables 1–3). Overall, SNP heritability on the liability scale was estimated at ∼28 % assuming the disease prevalence of 20–30%. Almost 40% of the variance explained by our top SNPs was attributable to the polymorphism in the *CASZ1* gene involved in blood vessel development. This strong signal has previously been revealed by “23andMe”^12^ and subsequently replicated in our own sample of ethnic Russian individuals^10^. An especially interesting finding, in our opinion, is a novel association of VVs with SNPs in a recently discovered *PIEZO1* gene. *PIEZO1* encodes a pressure-activated ion channel which senses shear stress and controls vascular architecture^27^.

Mice embryos lacking functional Piezo1 exhibit defects in vascular remodeling and die at midgestation^61^. Other prioritized genes were *PPP3R1, EBF1, STIM2*, and *HFE*. The latter gene has been linked to the risk of VVs in our recent candidate-gene study^19^. Meanwhile, we could not prioritize any gene in the locus tagged by rs3101725. On the one hand, this region contains two genes (*SLC12A2* and *FBN2*) that could play a role in varicose transformation, and on the other hand, the causal polymorphism was shown to be eQTL for a nearby non-protein coding RNA *LINC01184* with unknown function. The revealed genes can be considered as good candidates for future follow-up functional studies. It is noteworthy that *SLC12A2, FBN2, STIM2*, and *HFE* were included in the druggable gene set by Finan et al.^62^, and the *PIEZO1* gene product belongs to a “potential drug target” class according to the Human Protein Atlas (https://www.proteinatlas.org/).

Gene set enrichment analysis involving both genome-wide and less strongly associated signals detected gene categories related to abnormal vascular development and morphology. This observation is consistent with the hypothesis that pathological changes in the vein wall are the primary event preceding VVs formation^63^. Furthermore, our genetic correlations analysis confirmed known epidemiological associations between VVs and DVT^1^,^16^ as well as standing job^1^,^64^,^65^, rough labour^6^5,66, and lower levels of education^64^ (Fig.4, Supplementary table 11). Shared familial susceptibility with venous thromboembolism has already been shown by Zöller et al^67^. Here we demonstrate that DVT and VVs share specific genetic components which are independent from other factors such as obesity or number of operations (Supplementary fig. 3). Since none of the top GWAS hits were related to thrombosis, we can conclude that SNPs with less prominent associations are responsible for this genetic overlap. On the contrary, albeit several identified loci were also associated with blood pressure/hypertension and red blood cell traits, including mean corpuscular hemoglobin concentration (Table 2, Fig. 3), we found no evidence for genetic correlation between VVs and these traits. We can therefore attribute these effects only to pleiotropy. Intriguingly, we observed small, but significant genetic overlap with smoking (rg = 0.16). Smoking is considered only as suggestive risk factor for VVs since epidemiological studies have mainly shown no association with this habit^1^,^64^,^66^,^68^. Other novel interesting findings include genetic links with prospective memory and fluid intelligence (negative correlation), pain (knee pain, pain all over the body, neck or shoulder pain, and leg pain on walking), usual walking pace, and gonarthrosis.

Further, we obtained strong evidence for association between VVs and anthropometric traits such as weight, height, waist and hip circumference. We not only observed genetic correlations (Fig. 3, Fig. 4, Supplementary tables 10, 11), but also demonstrated these traits to be causative factors for VVs development (Fig. 5, Supplementary table 12). It is important to note that genetic overlap and causal relationships were inferred for both fat and fat-free mass. Thus, we can speculate that increased weight is a risk factor for VVs regardless of whether it was caused by excess body fat or a large mass of other tissues. According to some theories, association between overweight and VVs can be explained by greater concentrations of circulating estrogens or even by a confounding effect of parity^1^. Although this can be true, our results suggest that the cause may also be an increase in mass *per se*. Moreover, we found that height has common genetic component with VVs independent from weight and other traits (Supplementary fig. 4) and is also causally related to VVs (Fig. 5). Causal inference was additionally confirmed using an independent dataset (Supplementary table 13). This is in agreement with the results of the Edinburgh Vein Study that has shown a significant relationship between increasing height and VVs^64^. However, the majority of epidemiological studies to date have been focused only on BMI and did not consider height or other body characteristics. Notably, the formula for BMI contains the square of height in the denominator. It is possible that the impact of height underlies the inconsistency in the results, when some studies show a positive association between BMI and VVs, while others reveal no effect^1^. We therefore recommend that future epidemiological studies collect and analyze data on height and weight along with data on BMI.

Last but not least, we detected the causal effect of plasma levels of two proteins – MHC class I polypeptide-related sequence B protein (MICB) and CD209 antigen. Both molecules are involved in innate and adaptive immunity. MICB is a ligand for the activating receptor NKG2D present on the surface of natural killer and some other immune-related cells. CD209, also known as DC-SIGN, is a C-type lectin receptor expressed on dendritic cells and macrophages. In our study, Mendelian randomization analysis for CD209 was performed using two genetic instruments – rs505922 and rs8106657 identified by Suhre et al^69^. The first SNP is located in the *ABO* gene responsible for ABO blood group determination. Allele rs505922 C tags non-O group and is in linkage disequilibrium with the allele A of the neighboring SNP rs507666 (D’ = 1.00, r^2^ = 0.39), which was found to be the top variant associated with VVs in the “23andMe” GWAS^12^ This association was validated in our previous study using data from UK Biobank^10^. At the same time, rs505922 explains nearly 40% of the variability of the plasma CD209 concentration (our estimate based on the published data^69^) with the allele C being linked to higher levels of CD209. It is tempting to speculate that association between the *ABO* gene polymorphisms and VVs as well as between VVs and blood group A^70^ is mediated by this protein. We conducted SMR/HEIDI analysis for the locus containing these SNPs and demonstrated that both associations with VVs and CD209 were related to the same causal polymorphism (*P*_SMR_ = 3.3e-06, *P* HEIDI = 0.75), that supports our hypothesis. Nevertheless, the role of CD209 as well as MICB in VVs etiology needs further experimental confirmation. Besides this, it should be taken into account that the level of CD209 in plasma does not necessarily correlates with its level on the surface of the cells, and circulating CD209 may possess its own specific function.

## Methods

### Study sample, genotyping, and quality control

All cases and controls analyzed in our study were derived from UK Biobank^13^. Sociodemographic, physical, lifestyle, and health-related characteristics of UK Biobank participants have been reported elsewhere^15^. In brief, enrolled individuals were aged 40–69 years, were less likely to be obese, to smoke, to drink alcohol, and had fewer self-reported health conditions compared with the general population. All study participants provided written informed consent, and the study was approved by the North West Multi-centre Research Ethics Committee. Genotyping was performed using the Affymetrix UK BiLEVE and the Affymetrix UK Biobank Axiom arrays. Details on DNA extraction, genotyping, imputation, and quality control (QC) procedures have been reported previously^71^.

The discovery stage of our GWAS was based on genetic association data provided by the Neale Lab (http://www.nealelab.is/) for 337,199 QC positive individuals (6,958 patients diagnosed with “I83: VVs of lower extremities” and 330,241 control individuals without this diagnosis). Information on data processing is given on the Neale Lab website (http://www.nealelab.is/blog/2017/9/11/details-and-considerations-of-the-uk-biobank-gwas). In short, phenotypes were analyzed automatically by the PHEnome Scan ANalysis Tool. SNPs QC criteria included minor allele frequency (MAF) > 0.1%, the Hardy-Weinberg equilibrium *P* > 1.0e-10, and INFO score > 0.8. After filtration, 10,879,183 autosomal SNPs were left for the analysis. Per-individual QC procedure included removing non-white British and closely related individuals, patients with sex chromosome aneuploidies, and individuals who had withdrawn consent from the UK Biobank study. Associations were adjusted for sex and the first 10 principal components from the UK Biobank sample QC file. We also adjusted the data for body mass index and deep venous thrombosis using GWAS summary statistics for these traits downloaded from the same resource, and corrected the results for potential confounding using the intercept^72^ as a genomic control inflation factor λ. Details are given in Supplementary Methods.

A replication stage was performed by a reverse inverse-variance weighted meta-analysis that yielded genetic association data for 71,256 individuals (Fig. 1). For these purpose, we derived summary statistics for 408,455 UK Biobank participants (10,861 cases and 397,594 controls) from the Gene ATLAS database^14^ (http://geneatlas.roslin.ed.ac.uk/) and used these data as a “minuend” dataset. A detailed description of our approach with statistical processing is presented in Supplementary Methods, in the study of Deng et al.^18^, and in our manuscript deposited on bioRxiv^17^. Data provided by the Gene ATLAS project were obtained for a cohort of both related and unrelated individuals of white British descent. The associations were calculated using Mixed Linear Models with adjustment for age, age^2^, sex, array batch, UK Biobank Assessment Center, and the leading 20 genomic principal components as computed by UK Biobank. Details of this study including QC criteria have been reported previously^14^. In brief, they excluded individuals with missing or conflicting data, non-biallelic variants, and SNPs with MAF <0.01% and/or the Hardy-Weinberg equilibrium *P* < 1.0e-50. The Gene ATLAS contains information for over 30 million variants. In our analysis, we focused on 10,829,469 SNPs that overlap the Neale Lab SNP set.

### Conditional analysis

Regional association plots were generated with LocusZoom tool (http://locuszoom.org/) for regions within 250 kb from the lead SNP. Only replicated loci were analyzed. Conditional and joint (COJO) analysis was carried out by a summary statistics-based method described by Yang et al.^73^ Calculations were performed using the GCTA software^74^. LD matrix was computed with PLINK 1.9 software (https://www.cog-genomics.org/plink2) using 1000 Genomes data for 503 European individuals (http://www.internationalgenome.org/data/). We claimed the presence of one independent signal per locus if none of the polymorphisms except the lead SNP passed the significance threshold of *P* = 5.0e-08.

### Literature-based functional annotation

We used regional association plots to identify genes located within associated loci. These genes were further queried for potential involvement in the processes relevant to VVs pathogenesis. For each gene, we scanned an Online Catalog of Human Genes and Genetic Disorders (OMIM, https://www.omim.org/), the NCBI Gene (https://www.ncbi.nlm.nih.gov/gene), and the Pubmed database to inquire into their biological functions. Furthermore, we interrogated whether other hypothetical varicose-related traits were associated with these genes according to previously published GWAS. The Pubmed search was performed using the gene name as a keyword as well as combinations of the gene name and “GWAS”, or “genome-wide association study”, or “varicose”, or “venous”, or “vascular”. The information obtained was used for the literature-based gene prioritization.

Additionally, we used a PhenoScanner^75^ tool (http://www.phenoscanner.medschl.cam.ac.uk/phenoscanner) to make a list of complex traits associated at *P* < 5.0e-08 with lead SNPs or other polymorphisms in high LD (r^2^ ≥ 0.7) with lead variants. 1000 Genomes-derived polymorphisms served as proxies for the analysis.

### VEP analysis

For the VEP^51^ analysis, we used a set of SNPs within replicated regions associated with VVs at *P* ≤ T, where log_10_(T) = log_10_(*P*_min_) + 1, and *P*_min_ is a *P*-value for the strongest association per locus (±250 kb from the independent hit). Analysis was performed using software available online (https://www.ensembl.org/info/docs/tools/vep/index.html).

### SMR/HEIDI analysis

SMR/HEIDI^52^ approach was used to test for potential pleiotropic effects of identified loci on VVs and other complex traits including gene expression levels in certain tissues. SMR analysis provides evidence for pleiotropy, but is unable to define whether both traits are affected by the same underlying causal polymorphism. The latter is specified by a HEIDI test that distinguishes pleiotropy from linkage disequilibrium. Summary statistics for gene expression levels was obtained from Westra Blood eQTL^76^ (peripheral blood, http://cnsgenomics.com/software/smr/#eQTLsummarydata) and the GTEx^77^ database (44 tissues, https://gtexportal.org). VVs summary statistics was obtained from the Gene ATLAS database since it enables the maximum statistical power due to the greatest sample size. At the same time, VVs in this GWAS is completely correlated at a genetic level with VVs in our discovery dataset (rg = 1.00, Supplementary table 1). Summary statistics for other complex traits was derived from the GWAS-MAP database (see below).

Nominal *P* for SMR test was set at 2.63e-05 (0.05/1,898, where 1,898 is the total number of probes for the gene expression analysis) and 1.89e-06 (0.05/7*3,787, where 7 is the number of loci and 3,787 is the total number of non-VVs traits in GWAS-MAP). For HEIDI analysis in a gene expression study, we used a conservative threshold of *P* = 0.01 (*P* < 0.01 corresponds to the rejection of pleiotropy hypothesis). For HEIDI in a complex traits analysis, we implemented a less conservative threshold of *P* = 0.001 since the number of independent test was much higher. Details of data processing are given in Supplementary Methods.

### DEPICT analysis

DEPICT analysis was performed using DEPICT^53^ software version 1.1, release 194 (https://data.broadinstitute.org/mpg/depict/) with default parameters. GWAS summary statistics was obtained from the Gene ATLAS database. We employed DEPICT for both genome-wide significant SNPs (*P* < 5.0e-08) as well as for loci associated with VVs at *P* < 1.0e-05. As in previous analyzes, we defined locus as ±250 kb from the lead SNP. The major histocompatibility complex (MHC) region was eliminated. Significance threshold was set at FDR < 0.05.

### GWAS-MAP platform

GWAS-MAP platform integrates a database which was created to study cardiovascular diseases and contains summary-level GWAS results for 123 metabolomics traits, 1,206 circulating proteins, 2,419 complex traits from the UK Biobank, and 8 traits related to coronary artery disease, myocardial infarction and their risk factors. We additionally added 6 VVs-related traits analyzed in the present study (our discovery and replication datasets; the Gene ATLAS data for VVs; and the Neale Lab data for VVs, BMI, and DVT) as well as 33 traits from the Gene ATLAS database that we supposed to be biologically relevant to VVs. Description of all 3,795 traits is provided in Supplementary table 9. GWAS-MAP platform contains embedded software for LD Score regression^72^, 2SMR analysis (MR-Base package^58^), and our implementation of SMR/HEIDI analysis^52^. Further details are given in Supplementary methods.

### Genetic correlations

Genetic correlations (rg) between VVs and other complex traits were calculated using LDSC software (https://github.com/bulik/ldsc/). We applied a cross-trait LD Score regression technique as previously described by Bulik-Sullivan et al.^78^ This method requires only GWAS summary statistics and is not biased by a sample overlap. We analyzed 861 heritable non-VVs traits from the GWAS-MAP database. Only traits with a total sample size of ≥ 10,000 and ≥ 1 million SNPs tested were included in the analysis. GWAS summary statistics for VVs was obtained from the Gene ATLAS database. Statistical significance threshold was set at 1.16e-05 (0.01/861). For 25 traits that passed this threshold with |rg| ≥ 0.2, we calculated a matrix of genetic correlations. Partial genetic correlations were estimated using the inverse of the genetic correlation matrix. Significance threshold was set at 3.1e-05 (0.01/325, where 325 is the number of pairwise combinations). For partial genetic correlations between VVs, standing height, and weight, nominal *P* was set at 3.3e-03 (0.01/3). Clustering and visualization was performed by the “corrplot” package for the R language (basic “hclust” function). Further details are provided in Supplementary Methods.

### 2SMR analysis

Casual relationships between 2,221 non-VVs phenotypes (“exposures”) from the GWASMAP database and VVs (“outcome”) were assessed by 2-sample Mendelian randomization (2SMR) as previously described by the MR-Base collaboration^58^ (http://www.mrbase.org/). All the details of our protocol are given in Supplementary Methods. Binary traits with the number of cases or controls less than 1000 were not included in the study. GWAS summary statistics for VVs was obtained from the Gene ATLAS database. Analysis was performed on the GWASMAP platform. Two 2SMR approaches were used: an inverse variance weighted meta-analysis of Wald ratios (IVW) and MR-Egger regression. The nominal *P* was set at 1.1e-05 (0.05/2*2,221, where 2 is the number of approaches). For traits that passed either IVW or MREgger test, we performed the Steiger test^59^ for identifying the correct direction of effect, and conducted the robustness analysis (our own approach). Traits that passed all the analyses were then subjected to sensitivity tests^58^, that included assessing heterogeneity in causal effects amongst instruments, horizontal pleiotropy test (based on the intercept in MR Egger regression), leave-one-out analysis, and Funnel plots inspection. The nominal *P* for the Steiger test was set at 2.3e-05 (0.05/2,221), and for the robustness analysis, horizontal pleiotropy test, and heterogeneity analysis – at 1.3e-03 (0.05/38, where 38 is the number of traits that passed both 2SMR analysis and the Steiger test). Leave-one-out and Funnel plots were examined manually. Sensitivity analyses revealed the presence of horizontal pleiotropy for the majority of traits. To correct for this confounder, we omitted all instrumental variables associated with VVs with *P* < 0.01, and then repeated IVW 2SMR and sensitivity analyses. Additionally, for the resulting set of traits, we searched for independent GWAS in the MR-Base database^58^ and conducted confirmatory IVW 2SMR analysis (where appropriate) with the MR-Base default parameters. The nominal *P* was set at 0.013 (0.05/4, where 4 is the number of traits).

## Acknowledgments

The work of ASS was supported by the Russian Science Foundation [Project No 17–75-20223]. The work of YAT was supported by the Russian Ministry of Science and Education under the 5–100 Excellence Programme.

The work of SZS was supported by the Institute of Cytology and Genetics [Project No 0324–2018-0017].

The development of GWAS-MAP platform was supported by grants from the Russian Ministry of Science and Education under the 5–100 Excellence Programme, British Council’s Institutional Links Programme for Novosibirsk State University and University of Edinburgh (Project reference No IL4277322879), and by PolyKnomics BV. We thank Maatschap PolyOmica for providing free of charge access to computational services.

We express our gratitude to the Neale Lab and the Gene ATLAS projects for providing GWAS summary statistics for UK Biobank traits. We sincerely thank Yurii Aulchenko for valuable discussion. We gratefully acknowledge Denis Gorev, Eugene Pakhomov, and Anna Torgasheva who contributed to the development of the GWAS-MAP platform. We acknowledge Eugene Phakomov who provided computational support with SMR/HEIDI analysis. We also would like to thank Mariya Smetanina and Maxim Filipenko for obtaining funding.

## Author contributions

YAT and ASS conceived the study and provided its design. YAT carried out statistical analyses. ASS worked with literature sources. ASS and YAT interpreted all data and wrote the paper. TIS, SZS, and YAT contributed to the design and implementation of the GWAS-MAP database. SZS contributed to statistical analysis. All authors reviewed and approved the final version of the manuscript.

## Supplementary tables

**Supplementary table 1.** Genetic correlations between VVs in different datasets included in the analysis.

**Supplementary table 2.** Top loci associated with VVs of lower extremities.

**Supplementary table 3.** Results of conditional and joint analysis for replicated loci.

**Supplementary table 4.** Complex traits associated with the revealed hits (found in a PhenoScanner catalogue).

**Supplementary table 5.** Results of the VEP analysis.

**Supplementary table 6.** Results of SMR/HEIDI analysis. Searching for pleiotropic effects on VVs and gene expression.

**Supplementary table 7.** Results of DEPICT analysis for SNPs with *P* < 5e-08.

**Supplementary table 8.** Results of DEPICT analysis for SNPs with *P* < 1e-05.

**Supplementary table 9.** Description of traits included in the GWAS-MAP database.

**Supplementary table 10.** Results of SMR/HEIDI analysis. Searching for pleiotropic effects on VVs and other complex traits. Associations that passed both SMR and HEIDI analyses.

**Supplementary table 11.** Genetic correlations between VVs and other complex traits.

**Supplementary table 12.** Results of hypothesis-free 2SMR analysis.

**Supplementary table 13.** Results of 2SMR analysis performed on independent datasets.

**Supplementary table 14.** Trait abbreviations, full names of traits, and their sources (SMR/HEIDI).

**Supplementary table 15.** Trait abbreviations, full names of traits, and their sources (genetic correlations).

## Supplementary figure captions

**Supplementary figure 1.** Regional association plot of −log_10_ (*P*)for SNPs located at the distance of ≤ 250 kb from the index SNP. Color of circles indicates the strength of linkage disequilibrium with the lead SNP based on the squared correlation coefficient (r^2^). Blue line indicates recombination rate (cM/Mb). Genes are indicated as blue bars under the plot.

**Supplementary figure 2.** A quantile-quantile plot for observed vs. expected distribution of *P-* values for χ^2^ statistics.

**Supplementary figure 3.** Matrix of partial genetic correlations between VVs and a subset of non-collinear complex traits. Color depicts the sign and the absolute value of genetic correlation coefficient (rg). Combinations with statistically significant correlations are marked with points. Full names of traits as indicated in the Neale lab and the Gene ATLAS databases are given in Supplementary table 15.

**Supplementary figure 4.** Matrix of partial genetic correlations between VVs, standing height, and weight. Color depicts the sign and the absolute value of genetic correlation coefficient (rg). Combinations with statistically significant correlations are marked with points.

**Supplementary figures 5 –42.** Leave-one-out and Funnel plots before and after removal of instrumental variables associated with VVs with *P* < 0.01.

